# *FOXK2* amplification and overexpression promotes breast cancer development and chemoresistance

**DOI:** 10.1101/2023.05.28.542643

**Authors:** Yang Yu, Wen-Ming Cao, Feng Cheng, Zhongcheng Shi, Lili Han, Jin-Ling Yi, Edaise M da Silva, Higinio Dopeso, Hui Chen, Jianhua Yang, Xiaosong Wang, Chunchao Zhang, Hong Zhang

## Abstract

Activation of oncogenes through DNA amplification/overexpression plays an important role in cancer initiation and progression. Chromosome 17 has many cancer-associated genetic anomalies. This cytogenetic anomaly is strongly associated with poor prognosis of breast cancer. *FOXK2* gene is located on 17q25 and encodes a transcriptional factor with a forkhead DNA binding domain. By integrative analysis of public genomic datasets of breast cancers, we found that *FOXK2* is frequently amplified and overexpressed in breast cancers. FOXK2 overexpression in breast cancer patients is associated with poor overall survival. *FOXK2* knockdown significantly inhibits cell proliferation, invasion and metastasis, and anchorage-independent growth, as well as causes G0/G1 cell cycle arrest in breast cancer cells. Moreover, inhibition of FOXK2 expression sensitizes breast cancer cells to frontline anti-tumor chemotherapies. More importantly, co-overexpression of FOXK2 and PI3KCA with oncogenic mutations (E545K or H1047R) induces cellular transformation in non-tumorigenic MCF10A cells, suggesting that *FOXK2* is an oncogene in breast cancer and is involved in PI3KCA-driven tumorigenesis. Our study identified *CCNE2*, *PDK1*, and Estrogen receptor alpha (*ESR1*) as direct transcriptional targets of FOXK2 in MCF-7 cells. Blocking CCNE2- and PDK1-mediated signaling by using small molecule inhibitors has synergistic anti-tumor effects in breast cancer cells. Furthermore, FOXK2 inhibition by gene knockdown or inhibitors for its transcriptional targets (CCNE2 and PDK1) in combination with PI3KCA inhibitor, Alpelisib, showed synergistic anti-tumor effects on breast cancer cells with PI3KCA oncogenic mutations. In summary, we provide compelling evidence that FOXK2 plays an oncogenic role in breast tumorigenesis and targeting FOXK2-mediated pathways may be a potential therapeutic strategy in breast cancer.

## Introduction

Breast cancer (BC) is the most commonly diagnosed malignancy and the leading cause of cancer-related mortalities in women (1,2). Recent multi-omics studies improve the understanding of the mechanisms and disease stratification of BC, indicating that it is a group of heterogeneous diseases with various morphologies and clinical behaviors (3). In the past decades, the anti-hormonal and anti-Her2 targeted therapies on specific BC subsets contributed to the significantly improved long-term survival (4). Recently developed targeted therapy agents, such as anti-PI3KCA mutant inhibitors and anti-CDK4/6 inhibitors, also demonstrate clinical benefits (5,6). However, the innate and acquired resistance to the targeted or conventional cytotoxic therapy regimens eventually causes cancer relapse and cancer-related death (7). Therefore, this disease still poses a huge threat to public health, clearly demanding more specific tumor markers and potential targets for individualized therapies.

DNA amplification in oncogenes plays essential roles in cancer initiation and progression. Although a number of amplified oncogenes are identified in BCs, only a few have been demonstrated to be causally involved and therapeutically important (8–13). The long arm of chromosome 17 (17q) is a hotspot of cancer-associated genetic anomalies. Amplification of several genomic regions in 17q, such as 17q23 and 17q12 amplicons, is strongly associated with poor prognosis and is a significant predictor of relapse in BCs (14–16). These amplicons are typically discontinuous and complex in structure, suggesting that multiple oncogenes in this chromosomal segment may be co-selected during breast tumorigenesis.

*FOXK2* gene is located at 17q25.3 and encodes a transcription factor belonging to the Forkhead Box (FOX) family (17,18), which controls multiple biological processes (17,19). Evidence shows that FOXK2 promotes tumor cell proliferation and metastasis in some cancer types. FOXK2 is overexpressed in colorectal cancer and promotes the malignant phenotype (20,21). In hepatocellular carcinoma (HCC), FOXK2 overexpression results in enhanced cell growth and migration (22,23). Cancer-promoting functions were also reported in papillary thyroid cancer and ovarian cancer (24,25). Contradictorily, FOXK2 was reported to suppress ERα-positive BC cell proliferation by down-regulating the stability of ERα (26).

FOXK2 amplification is observed in all subtypes of BCs, especially in the luminal B subgroup (27). Here we explored the role of FOXK2 in BC tumorigenesis through integrative analysis of public genomic and proteomic datasets in BC patients or cell lines. Our study suggests that *FOXK2* promotes BC tumorigenesis and targeting FOXK2-mediated oncogenic pathways may be a potential therapeutic strategy for developing novel therapies of BCs.

## Materials and Methods

### Cell culture

Human BC cell lines MDA-MB-231, MCF-7, HCC1954, MDA-MB-361, and non-tumorigenic epithelial cell line MCF10A were purchased from the American Type Culture Collection (ATCC). HEK293T cell line was obtained from ATCC and used for lentivirus packaging. MDA-MB-231, MCF-7, MDA-MB-361, HEK293T cells were grown in DMEM containing 10% heat inactivated fetal bovine serum (Invitrogen), 100 units/ml Penicillin, and 100 µg/ml Streptomycin (Invitrogen). All cell lines were authenticated via short tandem repeat (STR) analysis in MD Anderson Cancer Center. Mycoplasma testing was performed by LookOut® Mycoplasma PCR Detection Kit (MP0035, Sigma-Aldrich).

### Antibodies and inhibitors

The antibody for FOXK2 (ab5298) was from Abcam Biotechnology company (Cambridge, United Kingdom). The β-Actin antibody (A1978) was from Sigma (St. Louis, MO) and mouse IgG (sc-2025) were from Santa Cruz Biotechnology (Dallas, TX). Doxorubicin (Dox, D1515), etoposide (VP16, E1383), 5-Fluorouracil (5FU, F6627), and Dichloroacetate (DCA) (347795) were from Sigma (St. Louis, MO). CDK2 inhibitors, Dinaciclib (HY-10492) and Alpelisib (BYL-719, HY-15244), were from MedChemExpress (Monmouth Junction, NJ).

### Fluorescence in situ hybridization

A bacterial artificial chromosome (BAC) containing the *FOXK2* locus at chromosome 17q25 (RP11-117P2) was purchased from BACPAC Resources Center (BPRC). It is fluorescently labelled with spectrum red (Vysis, Downers Grove, IL, USA), by a nick translation method. The labelled BAC probe was hybridized to metaphase spreads of MCF-7 and HCC1954 cell lines. The hybridized slides were counterstained with DAPI, and the images were captured using the Quips Pathvysion System (Applied Imaging, Santa Clara, CA, USA).

### *FOXK2* knockdown in BC cell lines

To knockdown *FOXK2* in BC cells, the cells were transduced with control and two independent shRNA lentivirus vectors specific for *FOXK2* using standard protocol. Briefly, HEK293T cells were seeded in the 10-cm dish with a concentration of 2.5×10 ^6^ for lentivirus generation. After 24 hours, cells were transfected with 6 µg shRNA vector, 2 µg psPAX2 (#12260, Addgene), and 2 µg pMD2.G (#12259, Addgene) plasmids by using lipofectamine 2000 (Life Technologies). The supernatant containing lentivirus was collected 48 hours later. BC cells were transduced with the lentivirus in the presence of polybrene (4 µg/ml) and selected with puromycin (Sigma-Aldrich) (0.5 to 2 µg/ml) for 5 days. The *FOXK2* target sequences are AACCCAGCTGGCCTTAACACT and AAGACAGAAGTCACACTTGAA.

### Immunoblotting

Cells were harvested and lysed using RIPA buffer containing 50 mM Tris-HCl (pH 7.4), 1% NP-40 (IGEPAL CA-630) (#I8896, Sigma-Aldrich), 0.25% sodium deoxycholate, 150 mM NaCl, 1 mM EDTA, 0.1 mM sodium orthovanadate, 0.5 mM PMSF, 1 mM DTT, 10 µg/mL leupeptin, 10 µg/mL aprotinin, 1 mM benzamidine. Total cell lysate (50 µg) was used for SDS -PAGE and transferred onto PVDF membranes (Millipore). After blocking with 5% non-fat milk, membranes were incubated with primary antibodies at 4 ℃ overnight. HRP conjugated secondary antibodies and ECL Western blotting Kit (GE Health) were used for signal detection.

### Cell proliferation assays

Cell proliferation assays were performed using the CCK-8 from Dojindo Laboratories (Rockville, MA, USA) following the manufacturer’s instructions. *FOXK2* gene knockdown and control cells were plated in 96-well flat-bottomed plates at a concentration of 1×10 ^4^ cells per well at 37 ℃ for different time periods. A mixture of 10 µl of CCK-8 and 190 μl media was added into each well and the cells were incubated for another one hour. The absorbance of each well was measured at 450 nm using a microplate reader. Each experiment was performed in triplicates. A student’s *t* test was used to determine the statistical significance.

To determine the effect of small molecular inhibitors on cell proliferation, 0.5 to 1.0×10 ^4^ cells were seeded in each well of a 96-well plate and incubated overnight. Compound doses were added to each well individually and cultured for 48 hours. Cell viability was determined using CCK-8 assay. The IC50 value of single agents was calculated using CompuSyn software based on the data from the cell viability assay (ComboSyn. Inc. Paramus. NJ2007).

### Wound-healing assay

*FOXK2*-knockdown and control MDA-MB-231 cells were seeded into 12-well plates at the concentration of 2×10 ^5^ per well and incubated overnight at 37 ℃. Cell layers were scraped in a straight line using a 200 µL pipette tip perpendicular to the bottom of the well. After scratching, cell monolayers were gently washed to remove detached cells, then replenished with fresh medium. Images on phase-contrast microscope were captured every 4 hours until cells migrated into the middle.

### Migration assays

Transwell migration assays were conducted on MDA-MB-231 cells following *FOXK2* gene-expression knockdown. Transwell chambers (8 μm pore) (Corning) were coated with 10 μg/mL fibronectin (Sigma-Aldrich) overnight at 4 ℃, washed in PBS, and rehydrated with serum-free media for 30 minutes at 37 ℃. Media were removed and cells (1 × 10^5^/chamber) were added to upper chambers with 15% FBS in lower chambers as a chemoattractant. Chambers were incubated for 3 hours at 37 ℃ and 5% CO_2_. After fixation cells were stained with crystal violet.

### Cell cycle analysis

*FOXK2* knockdown or control cells were harvested after trypsinization and washed with cold PBS buffer. Ice-cold 70% ethanol was used to fix the cells for 30 minutes at 4 ℃. The cells were incubated with PI (50 µg/ml) and RNase (100 µg/ml) solutions for 30 minutes at room temperature in dark room. The cell cycle distribution was determined by flow cytometry.

### Colony Formation Assay

The soft agar assay was performed under a standard protocol. Briefly, a 0.5% base agarose layer was prepared in a 6-well plate with a volume of 2 ml per well. Then, *FOXK2* knockdown BC cells were mixed with 0.3% upper agarose layer at a concentration of 1.0×10 ^4^ per well. Cells were incubated at 37 ℃ and 5% CO_2_ for 2 weeks and were stained with 500 µl of 4% formaldehyde and 0.005% crystal violet (C3886, Sigma) for 4 hours. Optical images were captured under a microscope and colony numbers were counted. Each assay has triplicates.

### Chromatin immunoprecipitation (ChIP) assays

ChIP assays were performed by using a Chromatin Immunoprecipitation (ChIP) kit (Upstate). Briefly, BC cell lines were cross-linked with 1% formaldehyde. Cells were then washed with ice-cold 1×PBS, lysed, and sonicated on ice to produce sheared soluble chromatin. The soluble chromatin was precleared with Protein A Plus agarose beads (sc-2001, Santa Cruz Biotechnology) at 4 ℃ for one hour and then incubated with anti-FOXK2 (ab5298; Abcam) antibody or with control mouse IgG at 4 ℃ overnight. The immune complexes were collected on protein A agarose and eluted. Cross-linking of immunoprecipitated chromatin complexes and of input controls were reversed by heating at 65 ℃ for 4 h, followed by proteinase K treatment. The purified DNA was analyzed by quantitative PCR. The primers for ChIP assays are the following, ESR1-CHIP-F: ACAGAGGAAGACTCTGTCTC; ESR1-CHIP-R: AAGACATCCCTTACTCTT CC; CCNE2-CHIP-F: TTGAAACCCACCCAAGTGAG; CCNE2-CHIP-R: TGAGGCAACAGTTAGCAAG; PDK1-CHIP-F: GAGAAACAAGCAATGGGTC; PDK1-CHIP-R: AGAGGACCT TCCTTGATATC.

### Synergy studies

A method developed by Chou and Talaly was used (28) to evaluate the synergistic effects of Dinaciclib combined with DCA, and Alpelisib combined with Dinaciclib or DCA. Dose-response curves and IC50 values for inhibitors were determined by proliferation assays. Equipotent molar ratios of combinational inhibitors were used to treat BC cells seeded in 96-well plates for 24 hours. Single inhibitor was used as control. The combination index (CI) was assessed and calculated by CompuSyn software. CI ˂ 1 indicates synergistic interaction, whereas CI ˃ 1 is antagonistic, and CI = 1 is considered additive effect.

### RNA-seq

The total RNAs isolated from control and *FOXK2* knockdown cells were first qualitatively assessed prior to the RNA-seq library construction. The RNA-seq library was sequenced on an Illumina HiSeq platform. Raw reads were adaptor trimmed and sequencing quality was evaluated with FastQC (version 0.11.2) software. Metrics including quality scores, sequence duplication, and adaptor content were used to decide whether more filtering is needed before genome mapping. Clean reads were mapped onto the reference genome (version: GRCh38.p13) with reads aligner, HISAT2 (v2.1.0). The mappable reads were assembled into transcripts or genes with a fast and efficient assembler, StringTie software (v1.3.5). Only coding genes were retained for further analysis.

To identify genes differentially expressed between *FOXK2* knockdown and control cells, we conducted differential gene expression (DGE) analysis by DESeq2 (v1.28.1), an R statistical package. Fold changes are log2 transformed (log2FoldChange) and adjusted p values (padj) are calculated by using the Benjamini-Hochberg procedure to control false discovery rate. Genes are significantly up-regulated if log2FoldChange >= 1 and padj < 0.05 in *FOXK2* knockdown cells, or significantly down-regulated if log2FoldChange <= -1 and padj < 0.05. With these criteria, differentially expressed genes are selected for further analysis.

## Results

### *FOXK2* is amplified and overexpressed in BCs

To examine the status of *FOXK2* located on Chromosome 17q25 in BC, we retrieved The Cancer Genome Atlas (TCGA) datasets, including 910 tumor samples and 981 normal controls. We found frequent genomic amplifications of *FOXK2* in BCs compared with normal controls (Figure 1A) in all subtypes classified by PAM50 (29), with the highest frequency seen in the LumB subtype (10.2%) and the lowest in the LumA subtype (1.7%). The amplification of *FOXK2* was further confirmed in MCF-7 and HCC1954 cells by FISH assay (Figure 1B). By mining the published datasets from TCGA (30), we found that *FOXK2* increased significantly in BC patients compared with the non-tumor breast tissues, suggesting a contribution to tumorigenesis in BC (Figure 1C). The relationship between *FOXK2* mRNA or protein abundance and copy number alterations was also examined in a Clinical Proteomic Tumor Analysis Consortium (CPTAC) BC cohort (27). The results showed that both FOXK2 mRNA and protein levels are positively correlated with copy number gains in BCs (Figure 1D, E). Immunoblotting revealed the up-regulation of *FOXK2* in BC cell lines compared with the benign breast epithelial cell line MCF10A (Figure 1F).

**Figure 1.**
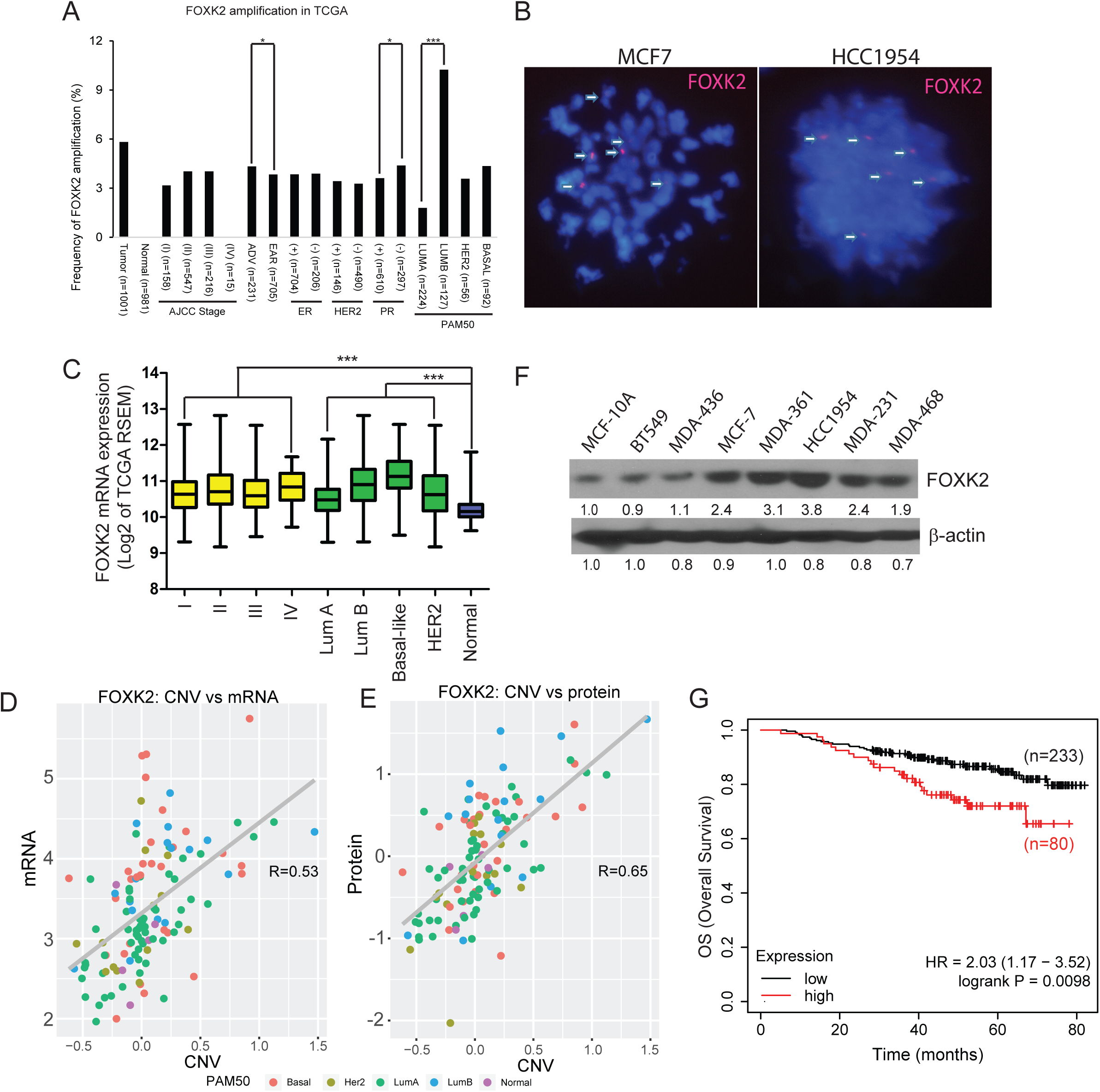
Integrative analyses of *FOXK2* in public genomic and proteomic breast cancer datasets. (A) *FOXK2* amplification across different subtypes of breast cancer. (B) FOXK2 amplification in MCF-7 and HCC1954 cell lines by FISH. (C) FOXK2 mRNA levels across different stages and subtypes of breast cancer. (D) Correlation between *FOXK2* copy number variations and its mRNA or (E) protein. (F) FOXK2 protein levels in multiple breast cancer cell lines by an immunoblotting assay. (G) Correlation of *FOXK2* gene expression with Overall Survival (OS) in untreated breast cancer patients by Kaplan-Meier survival analysis (http://kmplot.com/analysis/).

We further analyzed publicly available survival data from untreated BC patients through the Kaplan-Meier method (http://kmplot.com/analysis/). Comparing the *FOXK2* overexpressing group with the rest, *FOXK2* overexpression (measured by RNA-seq) was significantly associated with poor overall survival (OS) (Figure 1G), this correlation was also confirmed in BC subtypes (HER2+, HER2-/ER+, HER2-/ER-) (Figure S1).

### *FOXK2* knockdown inhibits the cell proliferation, anchorage-independent growth, cell cycle progression, and cell migration and invasion of BC cells

Effective knockdown of *FOXK2* by two lentivirus-mediated short hairpin RNA (shRNA) was confirmed by immunoblotting in four BC cell lines (MDA-MB-231, MCF-7, HCC1954, and MDA-MB-361) (Figure 2A). *FOXK2* knockdown resulted in a significant reduction in cell proliferation (Figure 2B). *FOXK2* knockdown led to less colony formation in four cell lines, suggesting FOXK2 is required for anchorage-independent growth of BC cells (Figure 2C, D). We next evaluated the effect of *FOXK2* knockdown on cell cycle distribution. As shown in Figure 2E, *FOXK2* knockdown in four BC cell lines resulted in a significant decrease in cell population in the S phase and an increase in the G0-G1 phase, indicating that *FOXK2* knockdown suppressed cell cycle progression.

**Figure 2.**
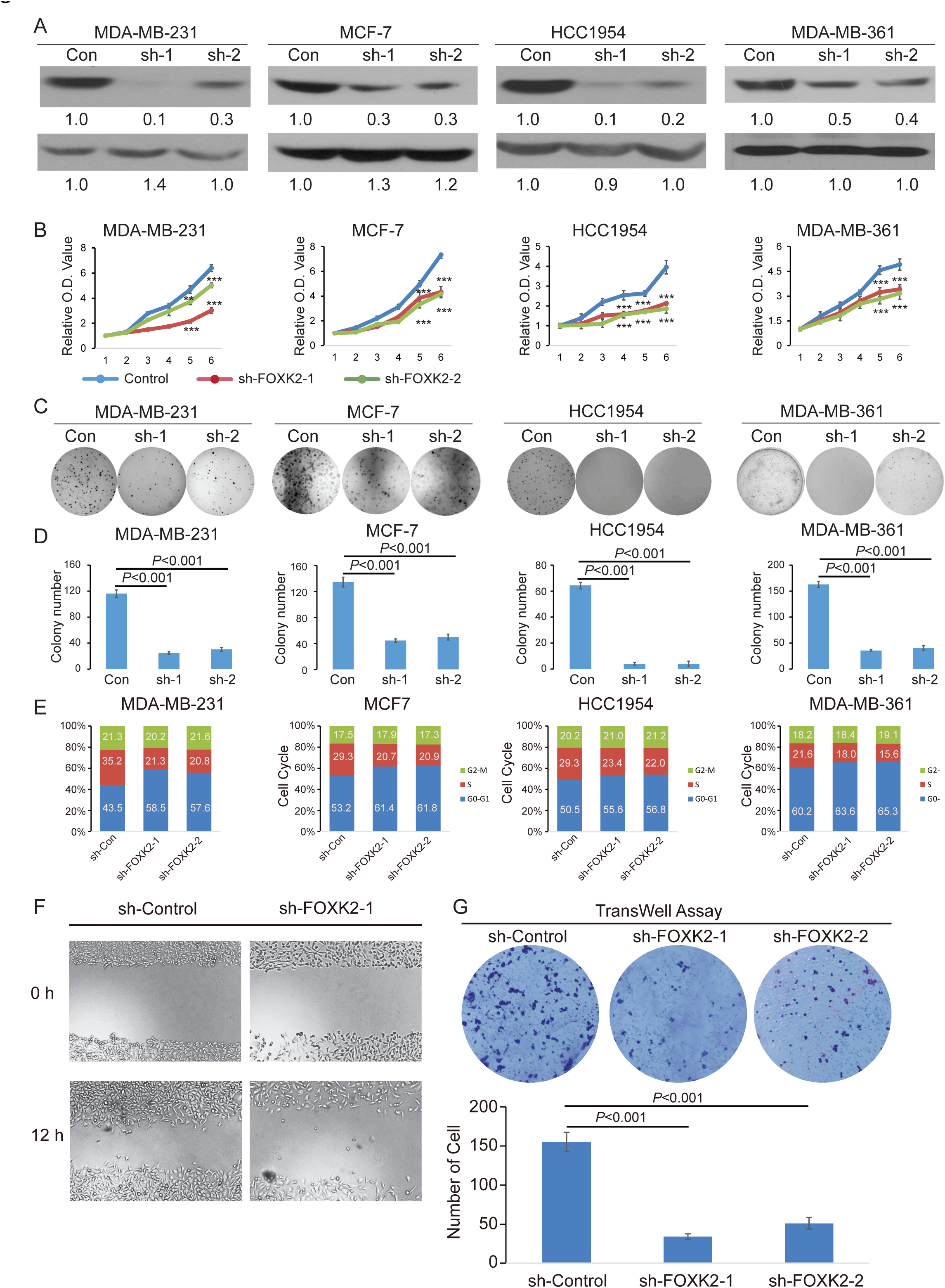
The effects of FOXK2 knockdown on cell proliferation, anchorage-independent growth, cell cycle progression, and cell migration and invasion of breast cancer cells in breast cancer cells. (A) *FOXK2* knockdown in MDA-MB-231, MCF-7, HCC1954, and MDA-MB-361 breast cancer cell lines by lentivirus-mediated expression of shRNAs. FOXK2 proteins are measured by an immunoblotting assay. (B) Cell proliferation in MDA-MB-231, MCF-7, HCC1954, and MDA-MB-361 cells determined by cck-8 assays following *FOXK2* knockdown. (C) An anchorage-independent growth (soft agar) assay of BC cell lines with or without *FOXK2* knockdown. (D) Mean colony numbers of four BC cell lines with or without *FOXK2* knockdown. (E) Cell cycle analysis by flow cytometry of four BC cell lines with or without *FOXK2* knockdown. (F) A wound-healing assay in which MDA-MB-231 cells with or without *FOXK2* knockdown. Confluent cell cultures were scratched, and wound closure was analyzed after 12 h. (G) A transwell invasion assay. Upper panel, cells that migrated across the porous membrane were stained with crystal violet. Lower panel, the number of cells with or without *FOXK2* knockdown.

Migration and invasion are crucial in spreading cancer cells, leading to local invasion and distant organ metastasis (31). To understand the functional relevance of FOXK2 in BC, we explored its role in cell motility and invasiveness. Firstly, we performed a wound-healing assay in *FOXK2* knockdown MDA-MB-231 cells. The gene knockdown significantly reduced the 2D migration ability of the MDA-MB-231 cells (Figure 2F). Additionally, we used Matrigel-coated Trans-Well chambers to examine the invasion properties of the MDA-MB-231 cells. Our results showed that *FOXK2* knockdown dramatically reduced the invasion of the MDA-MB-231 cells, suggesting a promotive role of FOXK2 in metastasis (Figure 2G).

To rule out off-target effects of the shRNA, we transfected BC cells with a *FOXK2* open reading frame (ORF) expression vector and then performed stable *FOXK2* knockdown with the sh-*FOXK2*-1 target sequence outside of ORF. The restored FOXK2 protein expression was resistant to sh-*FOXK2*-1-mediated knockdown (Figure S2A). As expected, this *FOXK2* expression reversed the proliferation defect (Figure S2B) and colony formation deficiency (Figure S2 C, D) caused by knockdown of endogenous *FOXK2*, further supporting that FOXK2 plays a critical role in BC cell proliferation and colony formation.

### C-terminal nuclear localization signal of FOXK2 plays a major role for its nuclear localization

By analyzing the transcriptome of BC cells, we found that the sequence ENST00000335255, encoding full length 660 residues, is the predominant transcript (Figure S3). FOXK2 protein has a forkhead-associated (FHA) domain, a FOX domain (FH), and two predicted nuclear localization signals (NLS) (Figure 3A). The NLS motif allows the active nuclear localization of the proteins. We examined the role of two NLSs in the nuclear localization of FOXK2. Wild-type FOXK2 (WT), N-Terminal NLS deleted FOXK2 (Mut-NLS-NT), C-terminal NLS deleted FOXK2 (Mut-NLS-CT), and both N- and C-Terminal NLS deleted FOXK2 (Mut-NLS-NT+CT) were fused with a green fluorescent protein (GFP) and transfected into HEK-293T cells. We found that N-terminal NLS deletion did not affect FOXK2 nuclear localization and C-terminal NLS deletion inhibited its nuclear localization, whereas dual deletion of both N- and C-terminal NLSs more severely inhibited its nuclear localization (Figure 3B,C). Using a rescue assay, we then examined the role of NLS of FOXK2 is required for its function in BC cell proliferation and found that the expression of FOXK2 with Mut-NLS-CT alone or Mut-NLS-NT+CT together failed to rescue *FOXK2* knockdown-mediated cell proliferation defect (Figure 3D). These results suggest that C-terminal NLS plays a major and N-terminal NLS plays an auxiliary role for FOXK2 nuclear localization.

**Figure 3.**
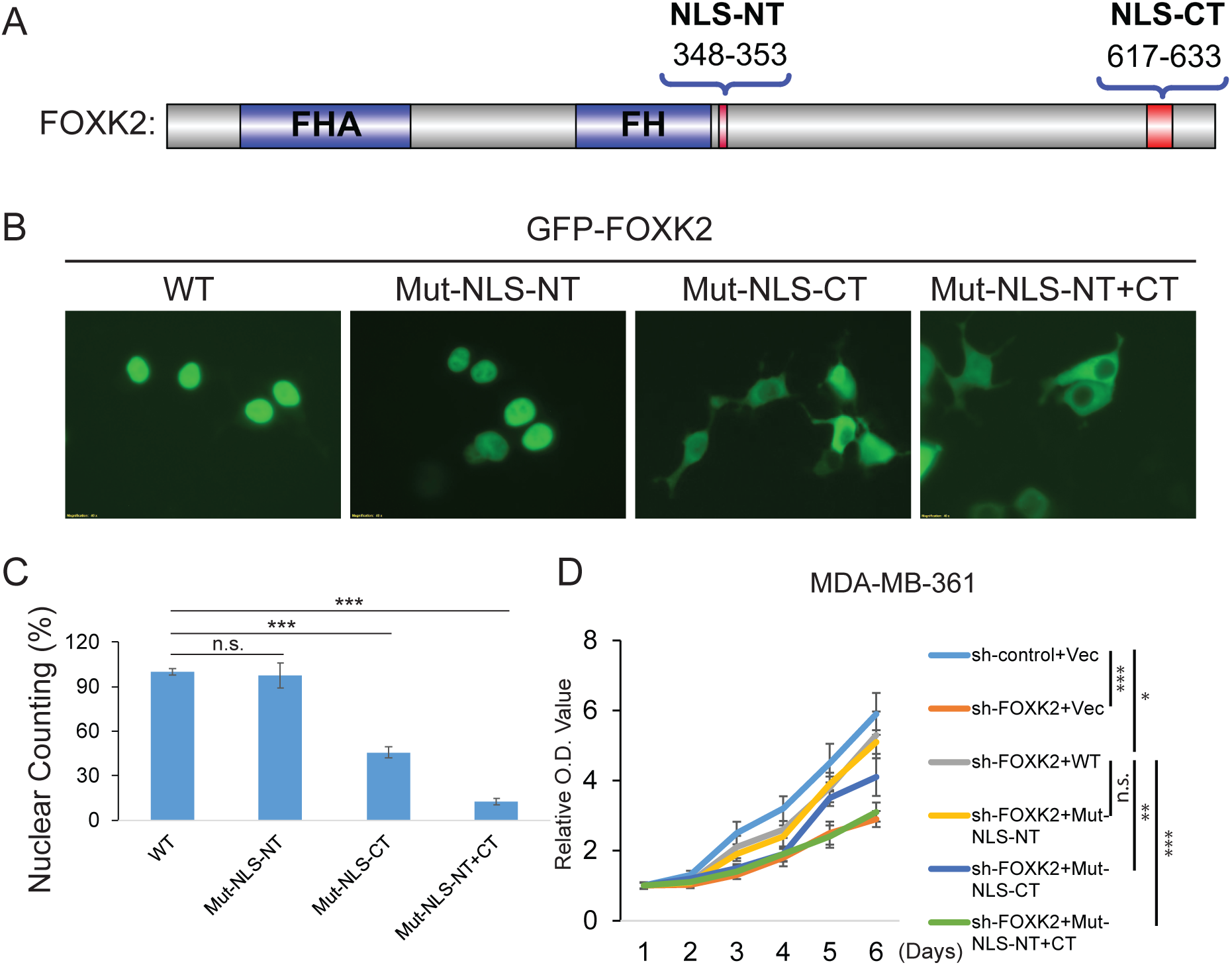
Determination of nuclear localization signal of FOXK2 protein. (A) Illustration of two domains in the FOXK2 protein sequence. (B, C) Determination of N-terminal NLS and C-terminal NLS domains in nuclear localization. (D) Mut-NLS-CT and Mut-NLS-NT+CT does not reverse the inhibition of cell proliferation in FOXK2 knockdown MDA-MB-361 cells.

### *FOXK2* drives MCF10A cell transformation by cooperating with RAS or PI3KCA oncogenic mutants

Our gene knockdown and rescue experiments suggest that FOXK2 is required for anchorage-independent growth of MDA-MB-231 cells, a basal B subtype with K-RAS mutation (32). To further examine the oncogenic role of FOXK2 in BC, we overexpressed FOXK2 or c-Myc, with oncogenic Ras in the non-malignant breast epithelial cell line MCF10A. Overexpression of FOXK2 or c-Myc alone did not result in MCF10A cell transformation, as determined by colony formation in soft agar. Co-overexpression of FOXK2 or c-Myc with K-RAS-V12D (a constitutively activated form of RAS) led to evident colony formation (Figure 4A, B). Notably, co-overexpression of FOXK2 and K-RAS-V12D resulted in fewer colonies than co-overexpression of c-Myc and K-RAS-V12D, suggesting that the transforming potential of FOXK2 may not be as strong as c-Myc.

**Figure 4.**
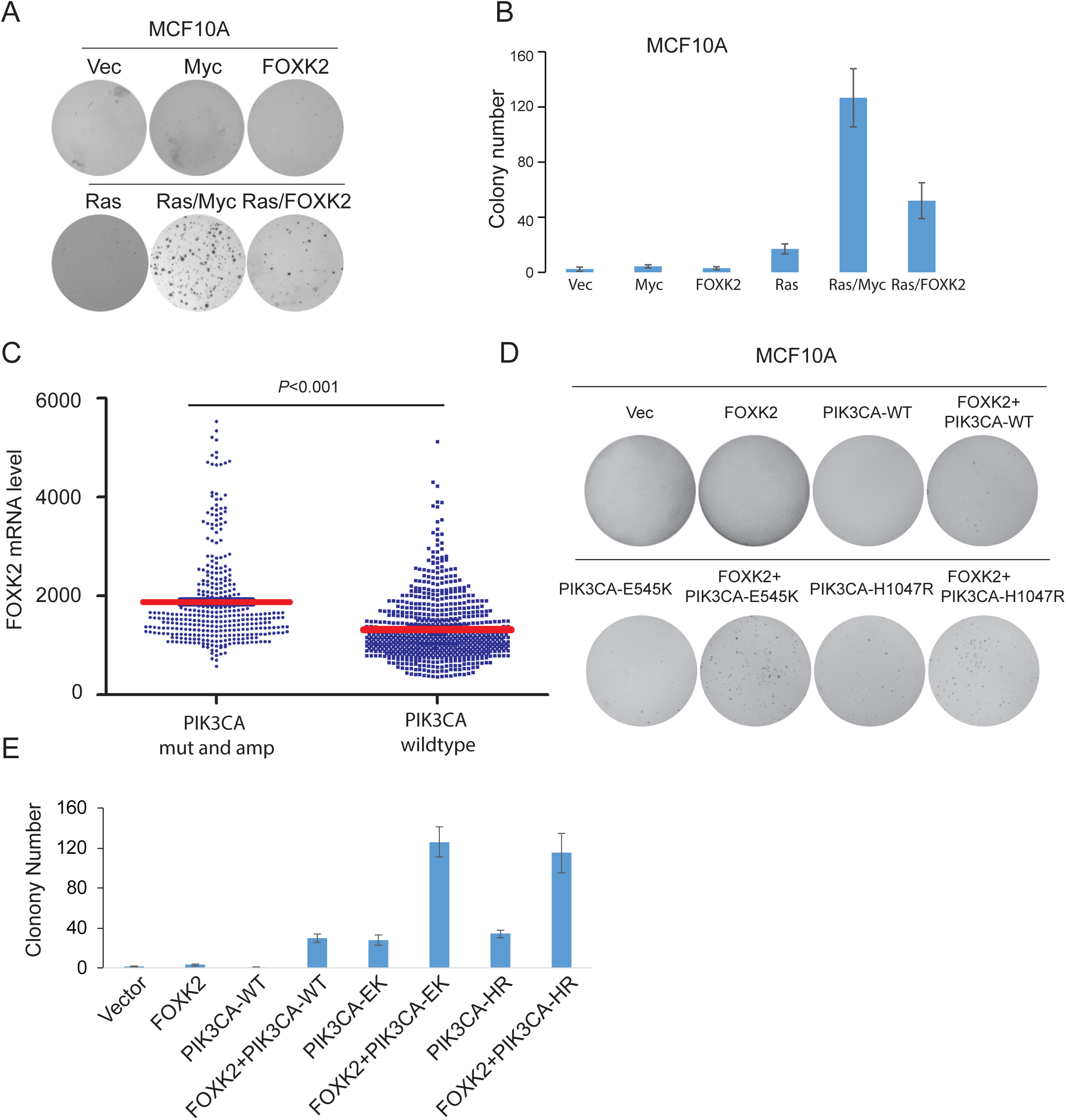
MCF10A cell transformation by *FOXK2* cooperatively with RAS or PI3KCA oncogenic mutants. (A) Anchorage-independent growth of MCF10A stably transfected with lenti-viruses expressing the indicated genes. (B) Quantification of colony numbers formed in each condition indicated in (A). (C) Comparison of *FOXK2* mRNA level in patient tumor samples carrying *PIK3CA* mutation/amplification with samples carrying wildtype *PIK3CA*. (D) Anchorage-independent growth of MCF10A overexpressing *FOXK2* and/or activating or wildtype *PI3KCA* alleles. (E) Quantification of colony numbers in each condition indicated in (D).

PIK3CA mutations are much more prevalent than RAS mutations in BCs, such as PIK3CA-E545K in MCF-7 and MDA-MB-361, and PIK3CA-H1047R in HCC1954. We investigated whether FOXK2 could cooperate with PIK3CA oncogenic mutants in BC tumorigenesis. First, we determined the correlation between FOXK2 expression and PIK3CA amplification and oncogenic mutations in BC patient tumor samples by analyzing TCGA data using the cBioportal platform (http://www.cbioportal.org/). *FOXK2* mRNA is higher in *PIK3CA* mutant/amplified samples compared with wildtype samples (*P* ˂ 0.001, student’s t-test) (Figure 4C). Next, we performed colony formation assays to evaluate the cellular transformation potential of FOXK2 in collaboration with PIK3CA oncogenic mutants in MCF10A cells. Overexpression of FOXK2 or PIK3CA wildtype alone in MCF10A cells did not lead to colony formation in soft agar. Co-overexpression of FOXK2 with PIK3CA mutants (E545K and H1047R) resulted in formation of many colonies, whereas overexpression of PIK3CA mutants alone or co-overexpression of FOXK2 with PIK3CA wild-type resulted in formation of a few colonies (Figure 4D, E). These results further support the oncogenic role of FOXK2 in BC and suggest it may cooperate with PIK3CA activating mutants to transform breast cells and promote BC progression.

### FOXK2 transcriptionally up-regulates *ESR1*, *PDK1,* and *CCNE2* in BC cells

To examine the FOXK2-mediated oncogenic signaling pathways and its target genes in BC, we conducted RNA-seq analysis on MCF-7 cells with *FOXK2* knockdown. We identified 609 down-regulated genes and 382 up-regulated genes in *FOXK2* knockdown MCF-7 cells (Figure 5A, Table S1). Gene set enrichment analysis of down-regulated genes by an over-representation test revealed inhibition of multiple pathways and their cross-talks (Figure 5B). Among them, *ERα(ESR1)*, *CCNE2,* and *PDK1* mRNA levels were significantly lower after *FOXK2* knockdown (Figure 5C). Since FOXK2 is a transcription factor, to determine whether *ESR1*, *CCNE2* and *PDK1* are direct targets of FOXK2, we performed a ChIP-qPCR assay to test whether FOXK2 binds to the promoter regions of these three genes in BC cells. Our results showed a high enrichment at the promoter regions of *ESR1* in MCF-7 and MDA-MB-361 but not MDA-MB-231 cells (Figure 5D). FOXK2 can also bind to the promoters of *CCNE2* and *PDK1* in all three cell lines tested (Figure 5D). We analyzed the expression of PDK1 and CCNE2 in normal breast tissues and invasive ductal breast carcinoma (IDBC) in the TCGA dataset using Oncomine (https://www.oncomine.org). The expression levels of PDK1 and CCNE2 are significantly higher in IDBC tissues compared with normal breast tissues (Figures 5E,F). The survival analyses by the Kaplan-Meier method revealed a worse OS in the groups overexpressing PDK1 and CCNE2 (Figures 5G,H). These findings indicate that *ESR1*, *CCNE2,* and *PDK1* are direct transcriptional targets of FOXK2 in BC cells and may mediate FOXK2 oncogenic signaling in BC.

**Figure 5.**
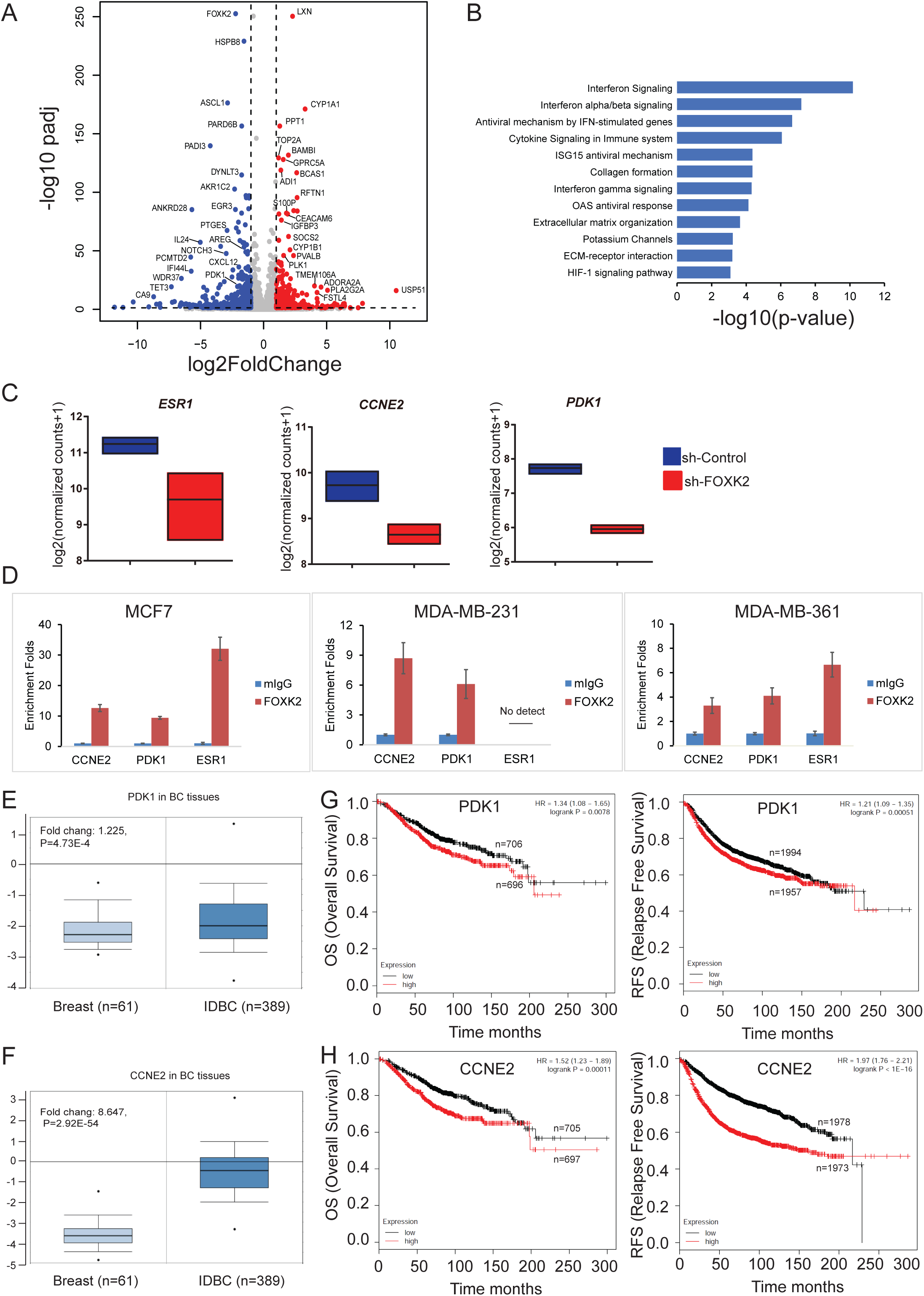
Identification of *FOXK2*-modulated genes in breast cancer cells. (A) Up-regulated (red dots) and down regulated genes (blue dots) in *FOXK2* knockdown MCF-7 cells by RNA-seq. (B) Gene set enrichment analysis of down-regulated genes by an over-representation test. (C) *ESR1 (ERα)*, *CCNE2* and *PDK1* were significantly decreased in *FOXK2* knockdown cells. (D) Determination of the binding sites of FOXK2 to *ESR1*, *CCNE2*, and *PDK1* promoter regions in MDA-MB-231 and MDA-MB-361 cells by ChIP real-time PCR. (E, F) The mRNA levels of *PDK1* and *CCNE2* in normal mammary tissues and invasive ductal breast cancer (IDBC) patient samples. (G, H) Association of PDK1 or CCNE2 overexpression with overall survival (OS) and relapse-free survival (RFS) by Kaplan-Meier survival analyses.

We next investigated FOXK2 highly correlated proteins in CPTAC luminal (LumA and LumB, ER-positive BCs) vs basal cases (predominantly triple-negative BCs). Proteins that are positively correlated with FOXK2 in luminal BCs (signed -log10FDR > 3) are mostly involved in translation processing (Figure S4 and Table S2), such as mitochondrial translational elongation and termination, translational elongation, and termination, or mitochondrial translation (GO terms). While those highly correlated proteins in basal cancers (signed -log10pval > 2) are mainly involved in mRNA processing, mRNA splicing, RNA splicing (Figure S4). In luminal cases, proteins that are negatively correlated with FOXK2 (signed -log10FDR < -3) are mostly related to protein transport, vesicle-mediated transport, and protein localization, and protein transport (Figure S4). A very few GO term enrichments were found in basal BCs. Our results revealed distinct subsets of FOXK2 modulated genes in different BC subgroups.

### Inhibiting FOXK2 or its related oncogenic signaling pathways may provide a potential therapeutic strategy for BC

We first evaluate the effect of *FOXK2* knockdown in response to conventional chemotherapeutic agents in BC cells. We treated FOXK2 knockdown and control cells with Doxorubicin, Fluorouracil (5FU), and VP-16. The results showed that *FOXK2* knockdown significantly enhanced the cytotoxic effect of the three chemotherapeutic agents in MDA-MB-231 cells (Figure S5A) and MDA-MB-361 cells (Figure S5B). These results show that inhibiting FOXK2 sensitizes FOXK2-overexpressed BC cells to conventional chemotherapeutic agents.

We hypothesize that FOXK2 oncogenic signaling pathway could be blocked by inhibiting its downstream target genes, *CCNE2* and *PDK1*. To test this, BC cell lines (MDA-MB-231, MDA-MB-361, and HCC1954) were treated with Dichloroacetate (DCA), a small molecule inhibitor of pyruvate dehydrogenase kinases (PDK) (33), or CDK2 inhibitor Dinaciclib (34,35). Our results showed that cell proliferation was inhibited in a dose-dependent manner in all three BC cell lines (Figures 6A,B). Using the method for synergistic analysis developed by Chou and Talaly (28), combination indexes (CIs) at the indicated combinations of DCA and Dinaciclib were derived from the cell lines tested using CompuSyn software (Figure 6C). DCA at constant centration of 5 mM could sensitize the chemotherapeutic response of Dinaciclib in tested BC cell lines (Figure 6C). We further examined the effect of DCA and dinaciclib, in combination with alpha-specific PI3K inhibitor Alpelisib, on BC cell proliferation. Alpelisib significantly suppressed cell proliferation in BC cell lines with activating PIK3CA mutation (MCF-7, HCC1954, and MDA-MB-361) compared with MDA-MB-231 cells which harbor wildtype PIK3CA (Figure 6D,E). *FOXK2* knockdown enhanced the cytotoxicity of Alpelisib in MCF-7 and HCC1954 cells (Figure 6F). Moreover, there were strong synergistic anti-cancer effects from DCA or Dinaciclib combined with Alpelisib in MCF-7 and HCC1954 cells (Figure 6G).

**Figure 6.**
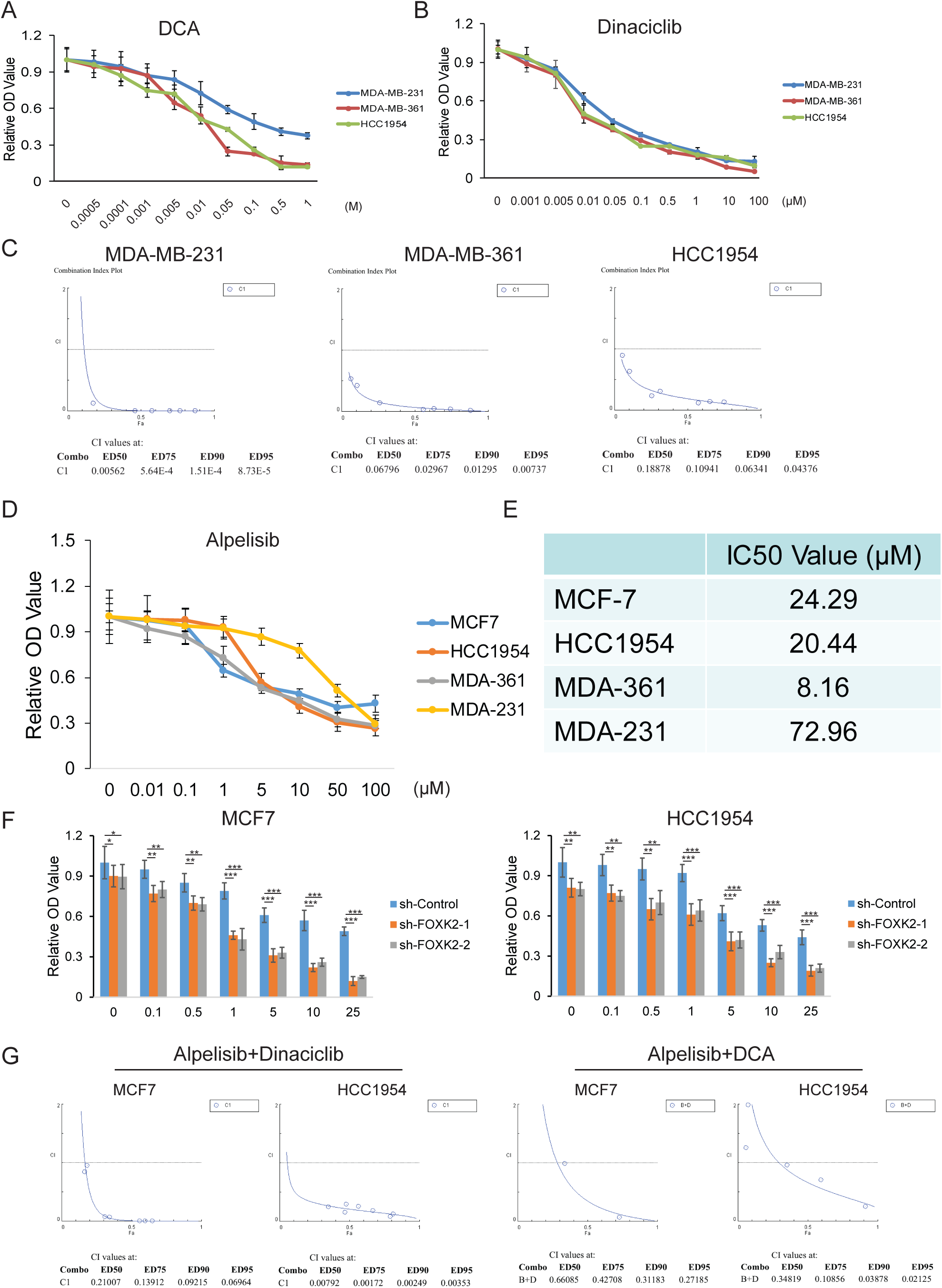
Synergistic anti-tumor effect of PDK1 inhibitor, CDK inhibitor, and PI3KCA inhibitor in breast cancer cell lines. (A, B) MDA-MB-231, MDA-MB-361, and HCC1954 cells were seeded into 96-well plates at a concentration of 1 ×10 ^4^ per well. After 24 h, cells were incubated with DCA (A), and Dina (B) for 48 h at indicated concentrations and cell proliferation was assessed by CCK-8 assay. (C) Synergistic anti-tumor effects of DCA with Dinaciclib in breast cancer cells. (D) MCF-7, MDA-MB-231, MDA-MB-361, and HCC1954 cells were treated with Alpelisib then cell proliferation was assessed by CCK-8 assay. (E) IC_50_ values of Alpelisib in four cell lines. (F) Cell proliferation by CCK-8 assays in MCF-7 and HCC1954 cells treated with Alpelisib and FOXK2 shRNAs. (G) Synergistic anti-tumor effects of Alpelisib with Dinaciclib or DCA.

## Discussion

Previous studies indicate that FOXK2 may be essential in regulating multiple signaling pathways, and its malfunction is implicated in severe developmental defects (36). However, the precise role of FOXK2 in tumorigenesis is under debate. Growing evidence shows that FOXK2 promotes tumor cell proliferation and metastasis in many types of cancers (21,22,24,25,37).

A few studies indicate an inhibitory role of FOXK2 in cancer progression by directly interacting with multiple transcription co-repressor complexes in BCs (38), especially in ER-positive breast tumors (26). However, the level of FOXK2 was reported to be higher in ERα-positive BCs than in ERα-negative counterparts and FOXK2 expression was positively correlated with ERα expression in ERα-positive BCs, making it questionable that FOXK2 plays a suppressive role in BCs. FOXK2 were also reported to inhibit cell proliferation in non-small cell lung cancer by down-regulating cyclin D1 and CDK4 (39). It is not clear whether FOXK2 function is highly context dependent and may vary in different cancer types.

Our proteogenomic analysis of FOXK2 in BC patients revealed that FOXK2 is positively correlated with cancer progression. *FOXK2* knockdown reduced cell proliferation, cell cycle progression and cell migration in multiple BC cell lines, unambiguously supporting an oncogenic role of FOXK2 in BCs. In addition, our data show that overexpression of FOXK2 with c-Myc or PI3KCA activating oncogenic mutations could transform mammalian epithelial cell MCF10A, suggesting that *FOXK2* amplification/overexpression and *RAS* or *PI3KCA* oncogenic mutation may cooperatively lead to cellular transformation and tumorigenesis in BC. Furthermore, we confirmed that the nuclear localization signal of *FOXK2* is required for its nuclear translocation and its cellular function in cell proliferation.

*ESR1* is overexpressed in 65%-80% of BCs and its high expression predicts clinical response to endocrine therapy (40,41). In this study we found that *ESR1* is a direct transcriptional target of FOXK2 in ER-positive BC cell line. This result suggests that FOXK2 plays its oncogenic role partially through up-regulating *ESR1* expression. Interestingly, in ER-negative MDA-MB-231 cells, FOXK2 was not found binding to the promoter region of *ESR1*. This suggests that the accessibility of the *ESR1* promoter region differs in ER-positive and ER-negative BC cells. Also, it suggests that FOXK2 regulatory network may be different according to the ER expression. Consistent with our RNA-seq data, the reported ER positively regulated genes, such as *PADI3*, *NDRG1*, *PTGES*, *CXCL12*, *SERPINE1,* and *THBS1,* are significantly decreased in MCF-7 cells with *FOXK2* knockdown (Figure S6). Interestingly, *CCNE2* and *PDK1* are direct transcriptional targets of FOXK2 in all BC cell lines tested. These results suggest that FOXK2-regulated gene networks may vary in different BC subtypes.

Previously, FOXK2 was claimed as a tumor suppressor and negatively regulated *EZH2* to inhibit tumor growth and metastasis of BC (38). Opposite to this claim, our data support an oncogenic role of *FOXK2* amplification and overexpression in BC. A positive correlation between *FOXK2* and *EZH2* was observed in studies by TCGA and Yu *et al*. (42) (Figure S7), suggesting that FOXK2 is not a *bona fide* negative regulator of *EZH2* in BC. FOXK2 may be oncogenic by up-regulating *ESR1*, *PDK1,* and *CCNE2* in BCs. Gene silence or inhibition specific to *ESR1*, *CCNE2* and *PDK1* suppresses cell proliferation in BC cells (43–45). TCGA BC datasets revealed that the expression levels of *CCNE2* and *PDK1* were significantly upregulated in tumor tissues compared with those in normal mammary tissues (Figures 5C,D). Our data clearly show that CCNE2 and PDK1 inhibitors can suppress the proliferation of BC cells, supporting the role of CCNE2 and PDK1 as downstream targets of FOXK2. Many CDK2 inhibitors are developed to inhibit the CDK2/cyclin E2 (an alias of CCNE2) complex activity since the biological functions of CCNE2 are predominantly dependent on CDK2 (46). Dinaciclib is a novel and potent inhibitor of CDK1, CDK2, CDK5, and CDK9, and its anti-tumor potency is being evaluated in various cancers (36,37). A phase II trial of dinaciclib in advanced BC suggests that dinaciclib monotherapy is not superior to capecitabine. Combined with other agents may be required for a better therapeutic effect (47). A phase II trial of DCA in metastatic BC was terminated due to higher risk and side effects (NCT01029925). As direct targeting transcription factors are challenging, to test whether anti-tumor potency could be achieved by inhibiting FOXK2-mediated kinases, we conducted cell cytotoxicity assays in three BC cell lines (MDA-MB-231, MDA-MB-361, and HCC1954) by using small molecular inhibitors (DCA and Dinaciclib) of PDK1 and CCNE2/CDK2 complex. Synergetic anti-tumor effects were observed by using DCA combined with Dinaciclib in MDA-MB-231 cells, which highlights a combination therapy of these small molecules targeting FOXK2-mediated kinases to treat FOXK2 amplified/overexpressed BCs. Furthermore, FOXK2 oncogenic signaling pathway inhibition with DCA or Dinaciclib showed synergistic anti-tumor effects with PI3KCA inhibitor Alpelisib in MCF7 and HCC1954 cell lines with PI3KCA activating mutations. Consistent with our study, PI3KCA inhibitor alpelisib has been reported to sensitize ER-positive BC cells to Tamoxifen, a selective modulator of ER function(48). These results suggest that FOXK2 inhibition can enhance sensitivity and overcome resistance of BC cells to current therapeutics.

In summary, we demonstrated that FOXK2 plays an oncogenic role in BC by transcriptionally activating ESR1/CCNE2/PDK1-mediated signaling pathways. Developing inhibitors targeting FOXK2 oncogenic signaling pathways may be a potential therapeutic strategy for BC.

## Acknowledgements

This research was funded by the Memorial Sloan Kettering Cancer Center institutional start-up fund (to HZ), and NIH-NINDS grant 5R01NS118008 (to JY). We also thank the technical support from the Cancer Prevention and Research Institute of Texas (CPRIT RP180734). Dr. Wen-Ming Cao’s present address is Department of Breast Medical Oncology, Zhejiang Cancer Hospital, Hangzhou, China.

## Author contribution statement

HZ and CZ designed the study. YY, WMC, LH, JLY, EMS and ZS carried out experiments. YY, HC, and CZ analyzed the data. JY, HD, CZ and CJ interpreted the results of the experiments. YY, CZ and FC drafted the manuscript. HZ, CZ, JY and XW edited the manuscript. All the authors approved the final version of the manuscript.

## Data availability statement

RNA-seq data are available in Gene Expression Omnibus (GEO) under accession number GSE230623.

## Supplementary Tables

**Supplementary Figure 1.**
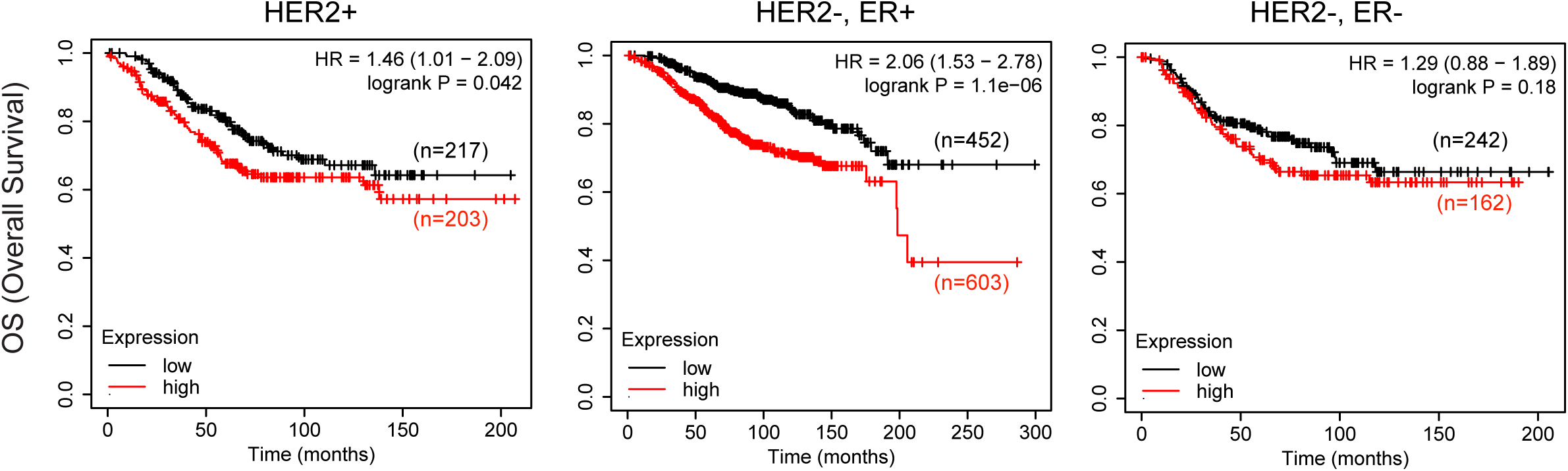
Correlation of *FOXK2* gene expression with Overall Survival (OS) in sub-groups of breast cancer patients.

**Supplementary Figure 2.**
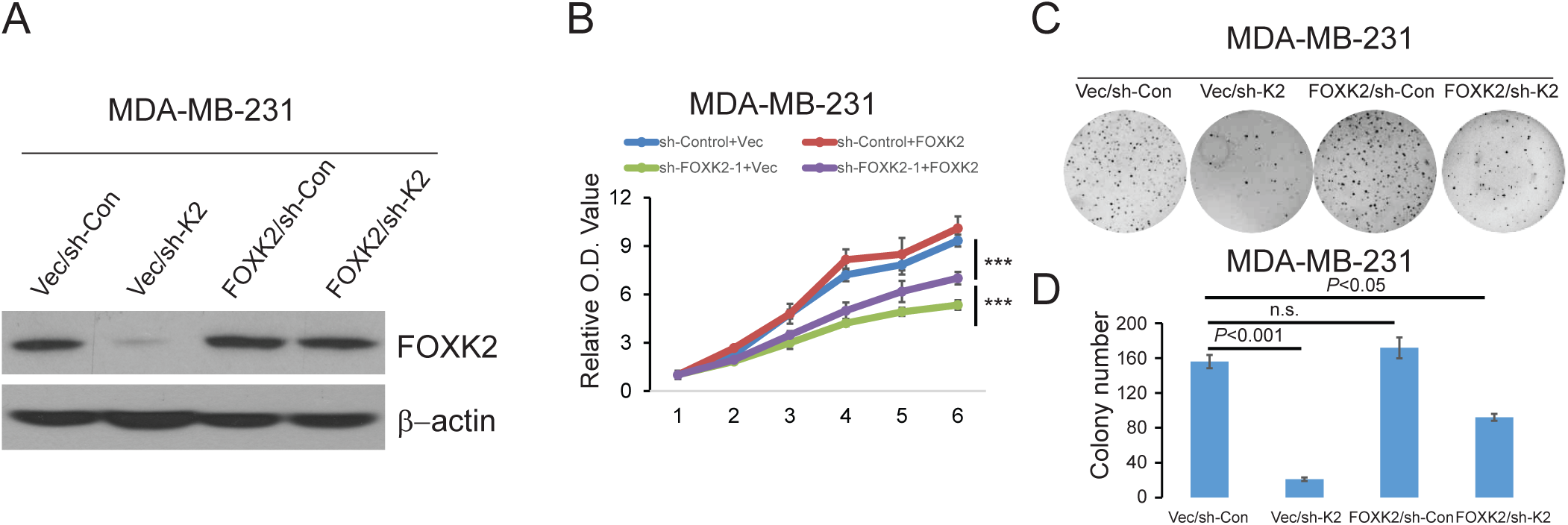
The specificity of *FOXK2* knockdown in breast cancer cell lines. (A) Overexpression of *FOXK2*-*ORF* to rescue *FOXK2* expression in MDA-MB-231 cells stably expressing *sh-FOXK2-1*. (B) Cell proliferation defect rescued by *FOXK2-ORF* in MDA-MB-231 cells. (C) Anchorage-independent growth defect rescued by *FOXK2-ORF* in MDA-MB-231 cells. (D) Mean colony numbers in MDA-MB-231 cells with different vectors. Error bars represent standard deviation from six samples. ** *P*<0.01, *** *P*<0.001, sh-FOXK2 versus sh-Control, t-test.

**Supplementary Figure 3.**
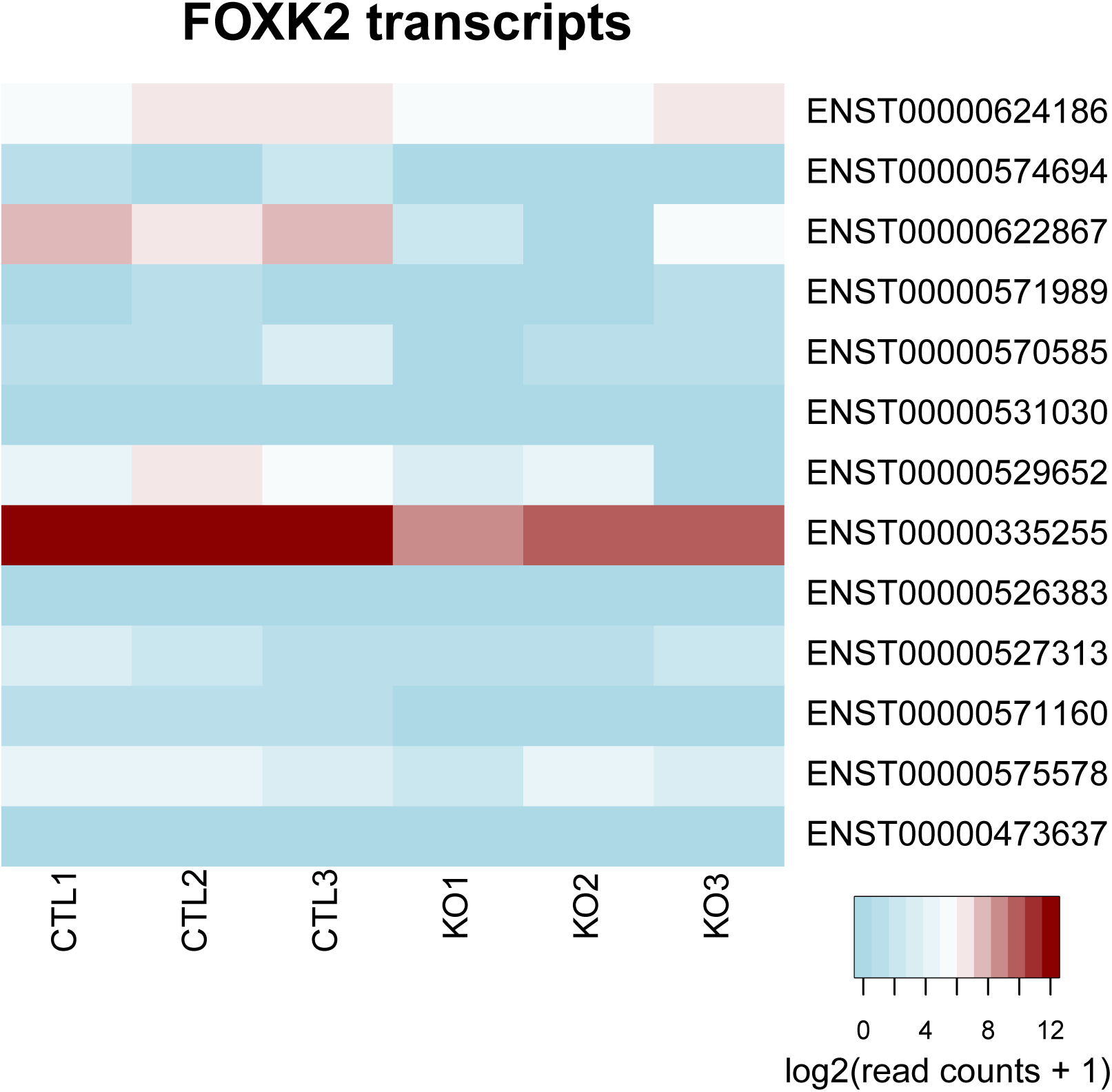
*FOXK2* transcripts in MCF-7 cells.

**Supplementary Figure 4.**
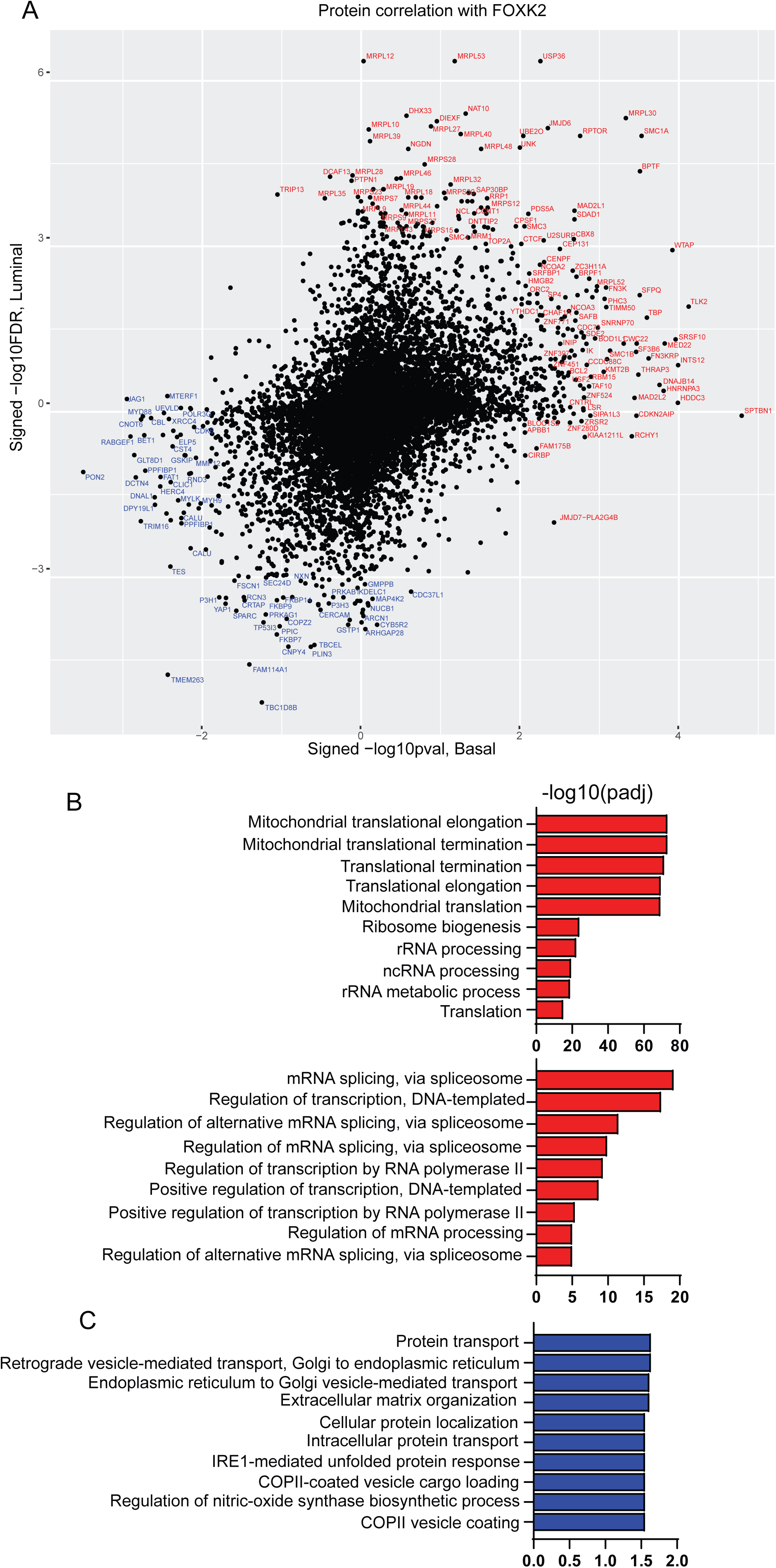
Proteins highly correlated with FOXK2 and their GO terms. (A) Pearson’s correlation of proteins with FOXK2 in PAM50 basal (x axis) and luminal (LumA and LumB, y axis) cases. Pearson’s correlation coefficients and their p values are calculated. P values are corrected by Benjamini-Hochberg procedure. Signed log10 p values (x axis, a relatively small sample size of basal cases) and signed log10 FDR-corrected p values (y axis) are plotted. (B) GO terms of proteins that are positively correlated with FOXK2 in luminal and basal cases. (C) GO terms of proteins that are negatively correlated with FOXK2 in luminal and basal cases.

**Supplementary Figure 5.**
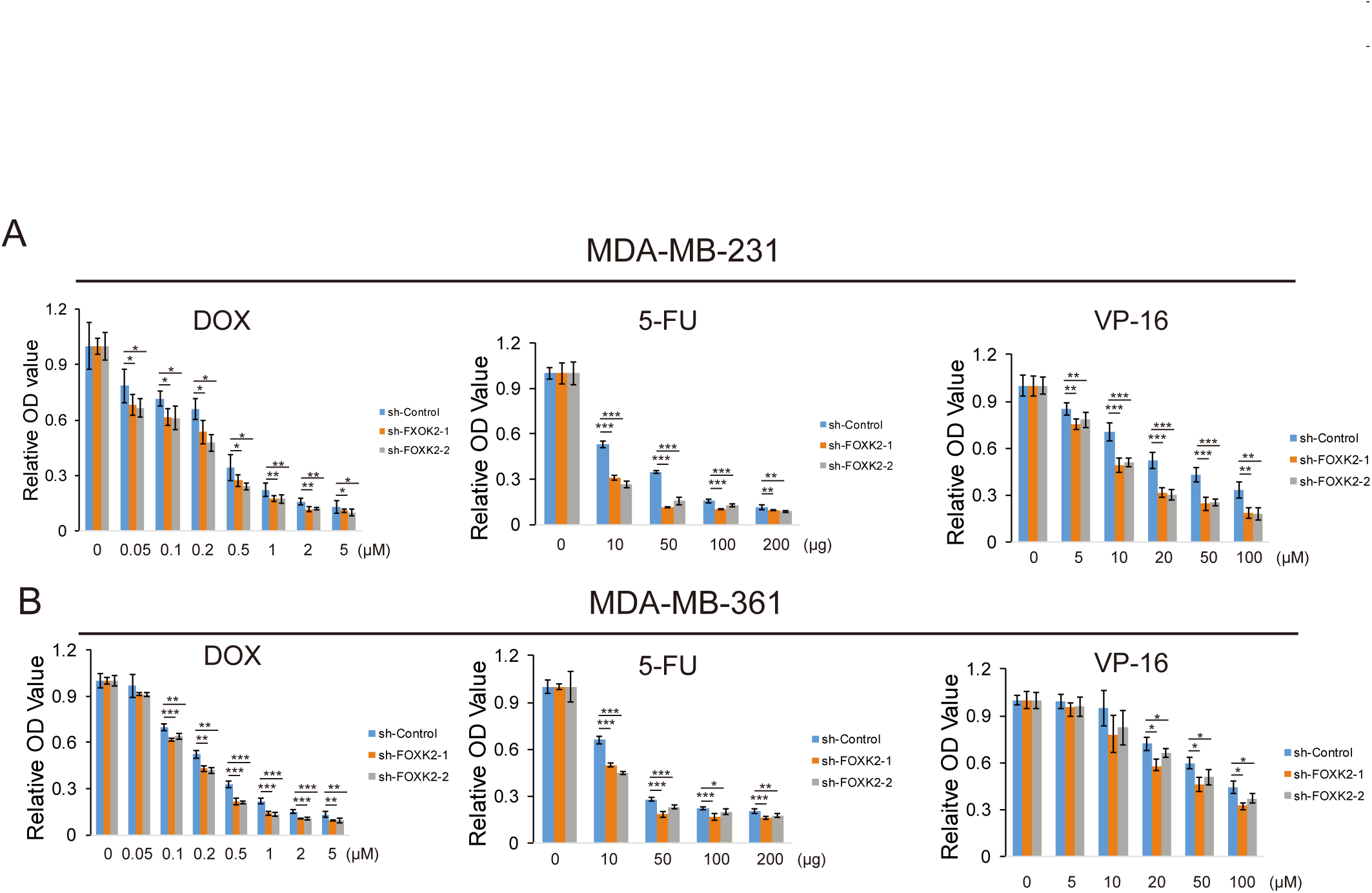
The effect of FOXK2 knockdown on the response to conventional chemotherapeutic agents in breast cancer cells. (A, B) *FOXK2* knocked down in MDA-MB-231 (A) and MDA-MB-361 (B) cells by two distinct sh-RNAs (sh-FOXK2-1 and sh-FOXK2-2). The knockdown cells were seeded into 96-well plates at a concentration of 1 ×10 ^4^ cells per well. After 24 hours, cells were incubated with Doxorubicin, 5-FU, and VP-16 for 24 hours at indicated concentrations and cell viability was assessed by CCK-8 assay. Results presented as % vehicle ± SD (n = 6). * *P*<0.05, \*\**P*<0.01, \*\*\**P*<0.001 (Student’s t test, two-tailed).

**Supplementary Figure 6.**
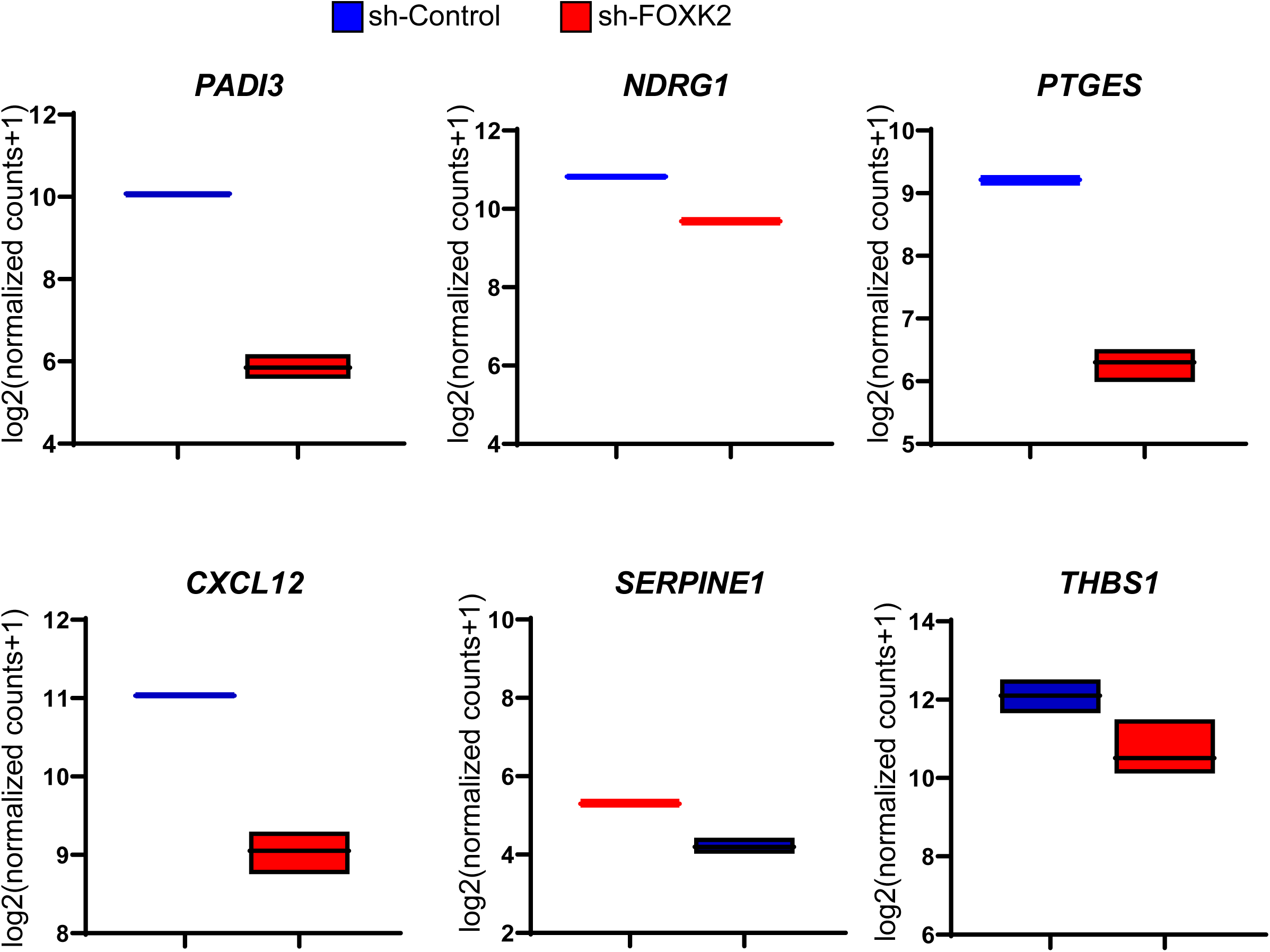
Expressions of ER targeted genes in FOXK2 knockdown MCF-7 cells.

**Supplementary Figure 7.**
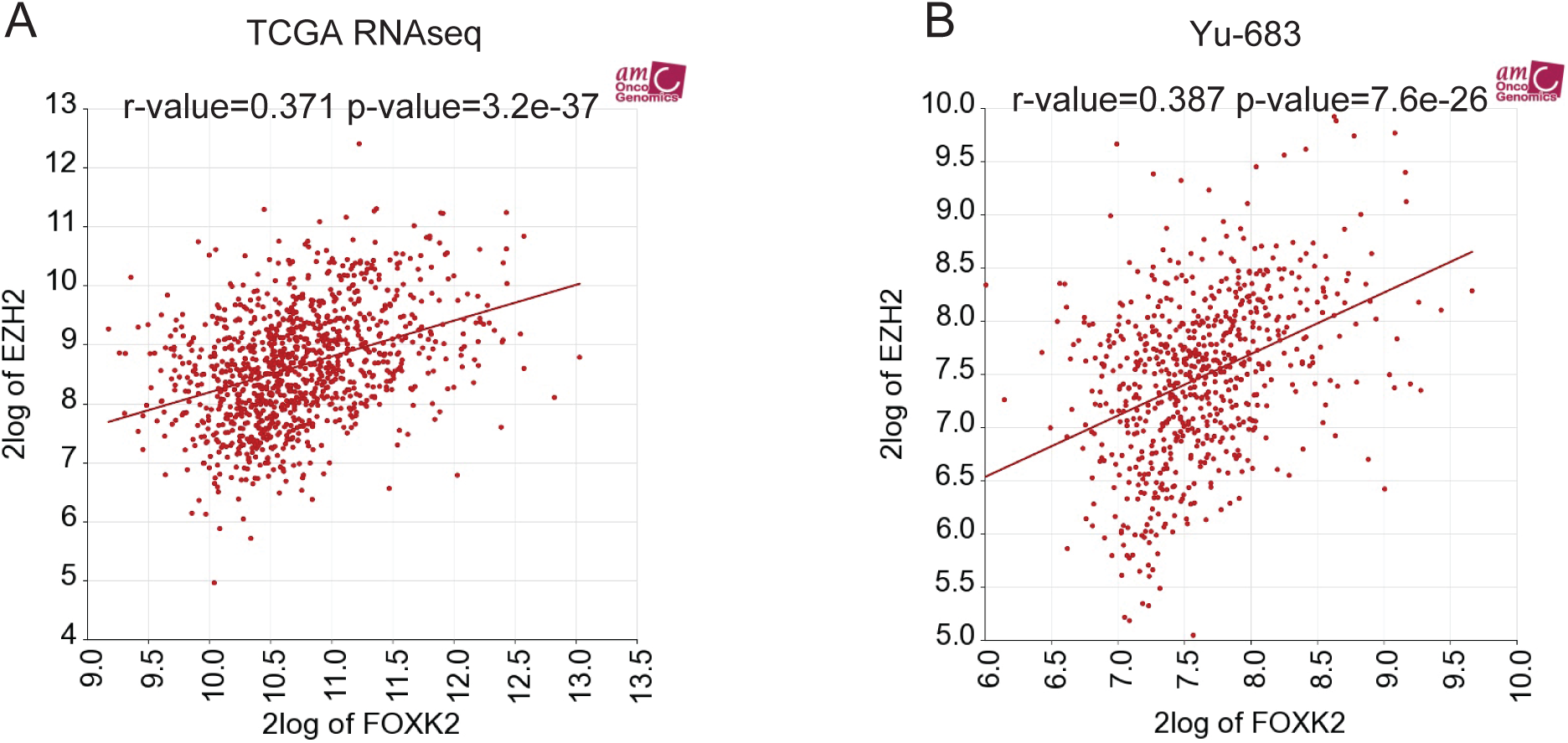
Correlation of FOXK2 and EZH2 in breast cancer tissues.

Table S1. Differentially expressed genes in FOXK2 knockdown MCF-7 cells.

Table S2. Pearson correlations between FOXK2 and other proteins in luminal and basal subtypes

